# Mastering Artifact Correction in Neuroimaging Analysis: A Retrospective Approach

**DOI:** 10.1101/2024.08.02.606374

**Authors:** Alícia Oliveira, Beatriz Cepa, Cláudia Brito, António Sousa

**Affiliations:** INESC TEC & University of Minho

## Abstract

The correction of artifacts in Magnetic Resonance Imaging (MRI) is increasingly relevant as voluntary and involuntary artifacts can hinder data acquisition. Reverting from corrupted to artifact-free images is a complex task. Deep Learning (DL) models have been employed to preserve data characteristics and to identify and correct those artifacts. We propose **MOANA**, a novel DL-based solution to correct artifacts in multi-contrast brain MRI scans. **MOANA** offers two models: the simulation and the correction models. The simulation model introduces perturbations similar to those occurring in an exam while preserving the original image as ground truth; this is required as publicly available datasets rarely have motion-corrupted images. It allows the addition of three types of artifacts with different degrees of severity. The DL-based correction model adds a fourth contrast to state-of-the-art solutions while improving the overall performance of the models. **MOANA** achieved the highest results in the FLAIR contrast, with a Structural Similarity Index Measure (SSIM) of 0.9803 and a Normalized Mutual Information (NMI) of 0.8030. With this, the **MOANA** model can correct large volumes of images in less time and adapt to different levels of artifact severity, allowing for better diagnosis.

## 1 Introduction

Magnetic Resonance Imaging (MRI) has become a crucial tool in medical diagnosis, providing relevant insights into the human body. Due to the inherent complexity associated with MRI procedures and their long examination times, susceptibility to movement increases, leading to image artifacts, particularly motion artifacts [15, 29], which result not only from patients’ voluntary movements but also from involuntary physiological movements (*e*.*g*., respiratory, cardiac and gastrointestinal peristaltic movements, vessel pulsations, blood and cerebrospinal fluid flow, and sudden involuntary movements such as swallowing) [29, 41]. Furthermore, other common artifacts in these exams are aliasing, magnetic susceptibility, and noise artifacts [35]. Among all the artifacts, motion-related abnormalities are particularly problematic, with a prevalence ranging from 10% to 42% in brain examinations [36].

Various strategies can be used to avoid motion artifacts in MRI, such as sedating patients, through moderate sedation or general anesthesia. However, these methods are not entirely effective, and motion artifacts are still present in around 12% of MRI images acquired under sedation and in around 0.7% of those acquired under general anesthesia [29, 41]. Given the limitations of existing preventive measures, correcting motion artifacts emerges as a viable solution to enhance medical imaging and improve diagnostic accuracy [12].

Advances in Deep Learning (DL) have shown great potential in correcting motion artifacts in MRI scans [14]. However, the other types of artifacts are not equally addressed. DL models can learn to identify and correct artifacts, leading to clearer and more reliable images for diagnosis. Artifact correction can be achieved through two primary approaches: prospective and retrospective. Prospective methods entail real-time correction during the MRI examination by modifying the data acquisition as it occurs [34]. Conversely, retrospective methods involve post-acquisition data enhancement after the MRI data acquisition is completed [34].

One key insight is that although real datasets present a high percentage of artifacts, publicly available ones are curated, lacking these and difficulting their usage for improving current correction models. Mimicking motion movements or any other artifact is of utmost importance when dealing with longitudinal cases. For instance, if one wants to correct a patient’s newly acquired MRI scan, a previous MRI scan without artifacts must be available to work as ground truth. Nonetheless, as these images are not publicly available, one must be able to emulate such artifacts.

Based on the necessity to recreate and correct artifacts, this paper proposes **MOANA**, a novel end-to-end retrospective method for correcting artifacts in multi-contrast brain MRI scans based on DL. Unlike previous solutions, **MOANA** is the first solution built to correct brain artifacts in four different contrasts while offering a simulation model that mimic several types of artifacts.

In brief, **MOANA** offers two models to enhance the pipeline of artifact correction. First, we offer a novel simulation model that can mimic motion, aliasing, magnetic susceptibility, and noise artifacts. Further, the second model corrects artifacts, emulated and real, by re-implementing the DL-based MC2-Net [28] model and optimizing and adapting it to a fourth contrast. As such, our approach can be applied to four types of contrast, including T1-weighted (T1w), T1-weighted contrast-enhanced (T1CE), T2-weighted (T2w), and Fluid Attenuated Inversion Recovery (FLAIR), in brain MRI scans, and can be adapted to other imaging modalities and medical applications, expanding the model’s potential in healthcare.

**MOANA**’s correction model has demonstrated remarkable performance in the different MRI contrasts. Specifically, in the T1w contrast, the model achieved Structural Similarity Index Measure (SSIM) [9] values of 0.9745 and a Normalized Mutual Information (NMI) [5] of 0.7811. In T1CE, it obtained an SSIM of 0.9607 and an NMI of 0.7929. In T2w, it attained an SSIM of 0.9792 and an NMI of 0.7821. Lastly, the **MOANA** model showed the highest performance in FLAIR contrast, with an SSIM value of 0.9803 and an NMI of 0.8030. Moreover, the model had significantly shorter training times, completing 500 epochs in approximately 22 hours.

The model enhances flexibility and adaptability by introducing a fourth contrast. It efficiently handles large volumes of images during training and testing, and has the ability to adjust to different levels of artifact severity, further enhancing its usefulness in improving image quality and facilitating accurate medical diagnosis.

## 2 Related Work

Küstner et al. [26] proposed automated methods for detecting and quantifying motion artifacts in MRI images. Their initial work utilized a Multi-Column Neural Network (MCNN) to detect motion artifacts in head and abdomen MRI images accurately. They further developed a retrospective correction method [27] using DL techniques, including Generative Adversarial Networks (GANs) [22] and Variational Autoencoders (VAEs) [25], which resulted in improved image quality metrics. Nonetheless, this approach may alter anatomical features, potentially affecting medical diagnosis.

Johnson and Drangova [24] introduced a method for motion correction by predicting artifact-free images from motion-corrupted data using a Conditional GAN (CGAN) [30]. While this method reduced motion artifacts, it also resulted in some irregularities and degradation of image details. Armanious et al. [12] implemented a retrospective method for correcting motion artifacts using Medical GAN (MedGAN) [13], which demonstrated high correction power. However, the loss of image details remains a limitation.

Lee et al. [28] proposed a method for correcting motion artifacts in multi-contrast brain MRI images, effectively reducing artifacts but showing limitations in correcting subtle motion artifacts. In [31], the authors established a technique for reducing motion artifacts using an Inception-ResNet [38] V2 DL architecture, showing promising results but with limitations in recognizing metallic implants. Tripathi et al. [39] modified the structure of CGAN [23] for motion artifact removal, achieving high accuracy but showing limitations in T1w contrast images. Finally, Singh et al. [33] presented a motion artifact correction model based on a Deep Neural Network (DNN) [17] and Hypernetwork [40], which produced sharp and accurate reconstructions that were particularly beneficial in real image scenarios.

**MOANA**’s correction model is a significant advancement when compared with these solutions, as it introduces a fourth contrast variation for correction, which extends the model’s flexibility and usefulness. Concerning flexibility, **MOANA** has the capacity to retain more information from larger quantities of data, improving its generalization for the four contrasts and the various types of artifacts, i.e., for each slice, the model leverages the information from each contrast and creates a more generalizable solution. This is also relevant for its usefulness, as this generalization can be exploited in real clinical scenarios, assisting in direct medical diagnosis.

## 3 MOANA

An overview of the **MOANA** model can be seen in Figure 1. The process includes data pre-processing, simulation of artifacts in the ground truth images, and correction of these artifacts using the correction model.

**Figure 1.**
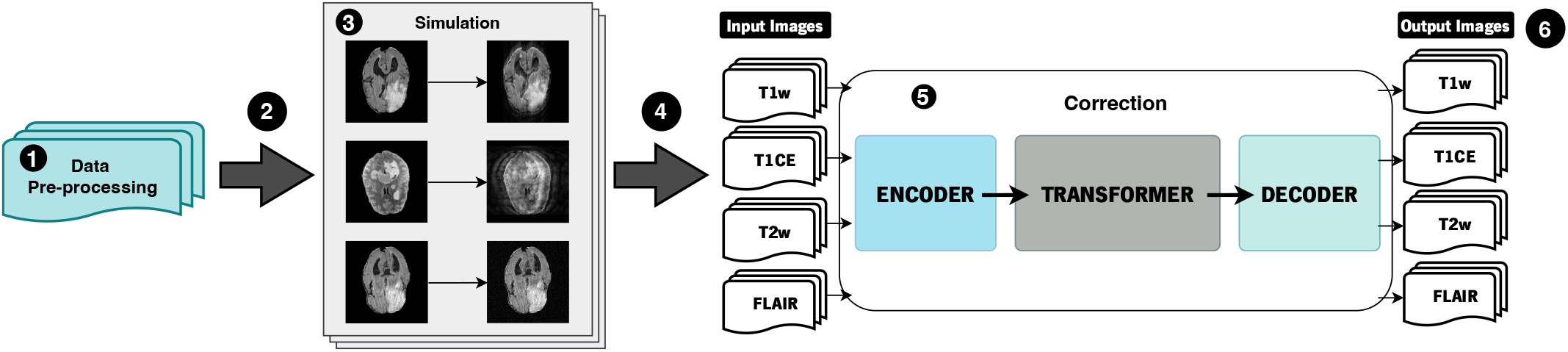
**MOANA**’s pipeline. The pipeline is composed of the following several stages: data pre-processing, simulation, and correction.

### 3.1 A day in the life of a corruption-free image

At the start of the workflow (Figure 1), input data is provided in the form of images that are free from artifacts. These images serve as the baseline for the subsequent processing steps. The input data is pre-processed (❶), involving tasks like rotation, resizing, and image selection. Once pre-processed, the data is fed into the simulation model (❷). During the simulation process, artifacts are introduced into the data. These artifacts result from different levels of artifact severity (❸). The corruption-free and the corresponding corrupted images are divided based on contrasts and introduced in the correction model (❹). This model is designed to analyze and correct any distortions or artifacts, thereby restoring the images to their original state, ideally in a corruption-free condition (❺). Finally, the corrected images serve as the output of the correction model. These images are now free from the artifacts and are ready for further medical analysis (❻).

### 3.2 Data Pre-processing

The pre-processing of medical images in the Neuroimaging Informatics Technology Initiative (NIfTI) format [18], as illustrated in Figure 2, comprises several sequential steps aimed at enhancing data quality and usability for the developed DL model. This process initiates with converting NIfTI volumes into the Portable Network Graphics (PNG) image format, which is compatible with the model, and is followed by a 90-degree rotation to optimize image orientation. Subsequently, specific brain slices for each image contrast are selected. Finally, the images are resized to 256*×*256 pixels to ensure uniformity and contribute to data normalization.

**Figure 2.**
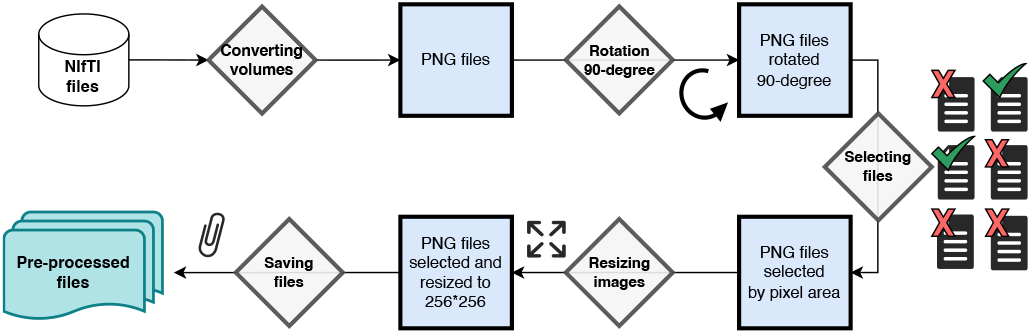
Pre-processing of medical images in NIfTI format involving conversion to PNG, rotation, selection of slices, and resizing.

### 3.3 Simulation Model

The simulation model is indispensable for enhancing the correction model due to the lack of publicly available datasets with corrupted images and the difficulty of obtaining real images (*i*.*e*., with artifacts) and the corresponding images without corruption (*i*.*e*., ground truth images). Thus, the simulation model enables the creation of artifacts from images without any corruption to obtain perturbations similar to real artifacts (*i*.*e*., motion, aliasing, magnetic susceptibility, and noise [35]). Our simulation model was based on the motion simulation model of [19] and was adapted to encompass different types of artifacts. These include simulating aliasing [32], reproducing fluctuations in magnetic susceptibility [16] and incorporating noise [21].

#### 3.3.1 Simulating Movement

The process of simulating movement in MRI images, based on a Fourier domain motion simulation model [19], starts by transforming the original image into the frequency domain using the Fast Fourier Transform (FFT) signal processing technique [20]. Subsequently, a trajectory of random movements is generated by adjusting the parameters explained below. This involves introducing random rotations and translations into the original image to create a corrupted version mimicking movement effects. The corrupted image is then transformed into the spatial domain using the Inverse Fast Fourier Transform (IFFT) [20].

Parameters such as standard deviation of rotations and translations are configured for simulating artifacts, with different ranges of *corruption scheme*.

#### 3.3.2 Simulating Aliasing

In this simulation, the original image is moved vertically by a specific distance, referred to as the *shift amount* [42]. This movement simulates the effect of undersampling. The moved image is then combined with the original image using a *blend factor*, which specifies the degree to which the shifted image influences the final image, thereby simulating the aliasing effect.

#### 3.3.3 Simulating Magnetic Susceptibility

This transformation is designed to simulate variations in magnetic susceptibility [16] in MRI images, caused by “the magnetic field inhomogeneity” [4] produced by materials (*e*.*g*., paramagnetic and diamagnetic materials). To simulate this artifact, a mask is applied to the source image with a specified *susceptibility factor*. This mimics the displacement effect seen with susceptibility differences.

#### 3.3.4 Simulating Noise

This alteration aims to introduce noise [21] into an image, controlled by a parameter termed *noise level*. To do this, a randomly generated noise matrix with the same shape as the original image is multiplied by the specified noise level and added to the original image.

The proposed simulation model considers two levels of artifact severity in medical images. At **level 1**, less severe artifacts are simulated, which include rotations and translations in the image (*i*.*e*., motion artifacts). At **level 2**, in addition to rotations and translations, the model increases the modifications introduced in the MRI images, simulating aliasing, magnetic susceptibility, and noise. The intention of the two levels is to create images with artifacts of different severity to improve the generalization of the correction model. The severity level of the simulation model depends on parameters such as the *shift factor*, the *susceptibility factor*, the *noise level*, and the *corruption scheme*. Figure 3 illustrates an example of each of these simulated artifacts.

**Figure 3.**
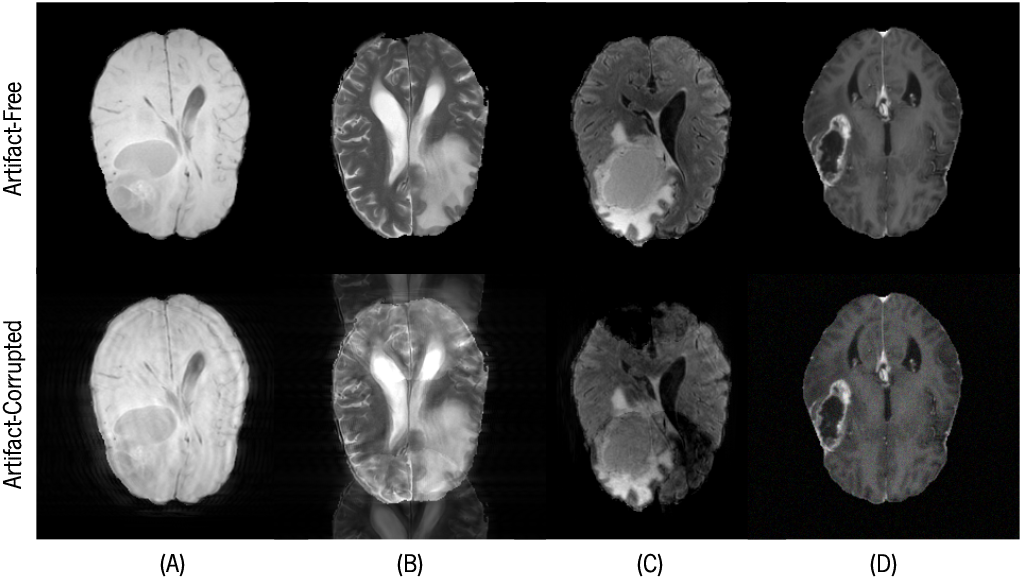
Examples of Simulation Model: (A) Simulating Motion in T1w contrast, (B) Simulating Aliasing in T2w contrast, (C) Simulating Magnetic Susceptibility in FLAIR contrast, and (D) Simulating Noise in T1CE contrast.

### 3.4 Correction Model

The proposed model, represented in Figure 4, is based on the MC2-Net [28] architecture, which is composed of a combination of two tasks: image alignment (including registration network and NMI maximization algorithm), and a motion correction network. Our model focuses exclusively on the correction model and aims to correct not only motion artifacts, but also aliasing, magnetic susceptibility, and noise attenuation. We have extended the capabilities of the original correction network architecture to support four contrast images, adjusting the input and output layers accordingly. Additionally, we have improved the model’s capacity by increasing the number of initial filters to 32, allowing for more feature extraction and better correction.

**Figure 4.**
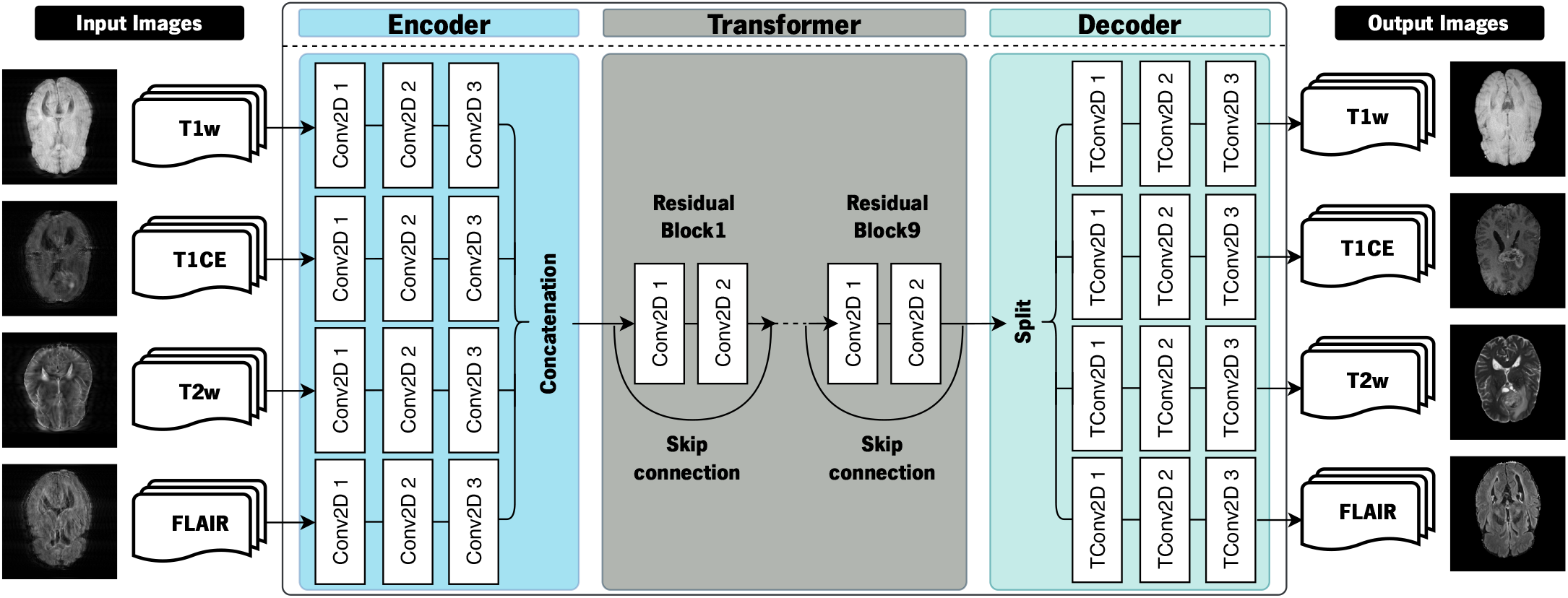
**MOANA**’s Correction Model Architecture. The DL-based model is composed of three main blocks, the *Encoder*, the *Transformer*, and the *Decoder*. Each of these blocks is composed of interconnected convolutional layers. The convolutional layer output is concatenated in the first block (Encoder). The Transformer is composed of 9 residual block layers and its output is split to serve as input for the Decoder.

The DL-based model is composed of three main blocks, the *Encoder*, the *Transformer*, and the *Decoder*. The Encoder transforms the input images into unique feature vectors for each contrast image, which uses 2D convolutions (*Conv2D*). These vectors are concatenated to allow the sharing of information between contrasts. The concatenated feature vectors are then passed through the Transformer, which comprises nine residual blocks, each containing two *Conv2D* interconnected by skip connections. The Decoder uses transposed 2D convolution (*TConv2D*) layers with Rectified Linear Unit (*ReLU*) activation to reconstruct the corrected features in the corresponding contrast image.

In order to train the model, we use a combination of loss functions that are customized for multi-contrast MRI correction. Specifically, we used SSIM [9] to ensure data consistency and VGG [11] perceptual loss to preserve perceptual details. We have chosen Adam [1] as the optimizer with a learning rate of 0.001. Our model has been implemented using TensorFlow 2.0 [10] and the Keras [3] library to ensure compatibility and ease of use.

#### 3.4.1 Image Quality Metrics

The correction network’s performance can be evaluated based on the quality of the resulting output image, using quantitative metrics [37]. With this, the proposed solution was assessed using three metrics: SSIM [9], NMI [6], and Normalized Root Mean Squared Error (NRMSE) [7].

SSIM [9] measures the structural similarity between two images by analyzing the loss of structural information. SSIM is defined as

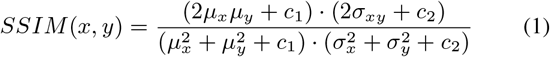

where “*µ*_*x*_ is the average value of *x, µ*_*y*_ is the average value of *y*, 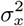 is the variance of *x, σ*^2^ is the variance of *y*, 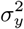 is the covariance of *x* and *y, c*_1_ = (0.01*L*)^2^ and *c*_2_ = (0.03*L*)^2^ are two variable to balance division with weak denominator, *L* is the dynamic range of pixel values” [39].

NMI [6] measures the agreement between the cluster labels assigned by two different clustering algorithms or methods by considering both the homogeneity within clusters and the separation between them [6]. NMI is expressed as

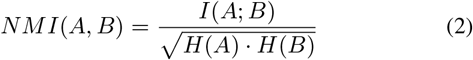

where *A* and *B* are the sets of clustering labels from two different clustering methods, *I*(*A*; *B*) is the mutual information between the clusterings, which measures the amount of information gained about one clustering from knowing the other, *H*(*A*) and *H*(*B*) are the entropies of the individual clusterings, which measure the uncertainty or disorder within each clustering [6].

The NRMSE [7] is a metric commonly used to assess the accuracy of a predictive model, regression model, or forecast. It normalizes the RMSE (Root Mean Squared Error) [8] by the range of the observed values [7]. NRMSE referred to as

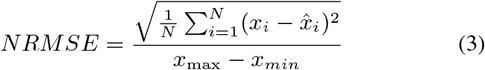

where *N* is the number of observations, *x*_*i*_ represents the actual observed value, 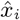 represents the predicted value, and *x*_max_ and *x*_min_ denote the maximum and minimum observed values, respectively [7].

## 4 Evaluation

We evaluate the effectiveness of **MOANA** artifact correction in terms of different levels of severity with different datasets. Our key insights are as follows:

- **MOANA** enhances state-of-the-art results when compared to the baseline model MC2-Net.
- **MOANA** allows the efficient addition of a fourth contrast for correction.
- **MOANA** provides different levels of severity regarding the simulation model, being able to attain results up to **0.9964** in SSIM for the highest severity (*i*.*e*., heavily corrupted images).

### 4.1 Experimental Setup

#### Testbed Setup

The tests were performed in two nodes, one with an NVIDIA GeForce RTX 2080 Ti (Node 1) and a second one with an NVIDIA GeForce RTX 3090 (Node 2). The training process is done by optimizing the correction network for several epochs, with hyperparameters tuned for optimal performance.

#### Workloads and metrics

The model was trained using data from the Brain Tumor Segmentation Challenge 2021 (BraTS 2021) [2]. This dataset consists of multi-parametric MRI (mpMRI) scans available in the Neuroimaging Informatics Technology Initiative (NIfTI) format (.nii.gz) that comprise native T1w, post-contrast T1w, T2w, and FLAIR volumes [2].

Four different subsets were created from the BraTS dataset: A1, A2, B1, and C1. Dataset A1 contains 48 subjects and 786 slices (262 slices from each contrast) selected from three contrasts (T1w, T2w, and FLAIR). Of these, 193 slices were used for training, 34 for validation, and 35 for testing. Dataset A2 includes 48 subjects and 1048 slices (262 slices from each contrast) chosen from four contrasts (T1w, T1CE, T2w, and FLAIR). The distribution of slices for training, validation, and testing follows the one of dataset A1. Dataset A1 was simulated with artifact severity levels 1 and 2, while dataset A2 was simulated with level 2. Dataset B1 consists of 3600 slices from 248 subjects, where each contrast has 900 slices for a total of four contrasts (T1w, T1CE, T2w, and FLAIR). The slices were split into three sets: 630 for training, 135 for validation, and 135 for testing. Dataset B1 was simulated with artifact severity levels 1 and 2. The last subset of the BraTS, dataset C1, consists of 140 slices (35 slices for each contrast) from 4 subjects. These slices contain real movement and were used to conduct real tests with artifacts (*i*.*e*., for model inference), evaluating the model’s capability to correct real artifacts.

Table 1 summarizes our datasets. Node 1 was used to train the model with datasets A1 and A2, and Node 2 was used with dataset B1. The model’s performance is validated by resorting to SSIM, NMI, and NRMSE metrics.

**Table 1.** Datasets used in our evaluation. The number of slices used for each phase (i.e., train, validation, and test) is provided per contrast.

|  | Subjects | Slices | Contrasts | Train | Validation | Test |
| --- | --- | --- | --- | --- | --- | --- |
| A1 | 48 | 786 | 3 | 193 | 34 | 35 |
| A2 | 48 | 1048 | 4 | 193 | 34 | 35 |
| B1 | 248 | 3600 | 4 | 630 | 135 | 135 |
| C1 | 4 | 140 | 4 | – | – | 35 |

### 4.2 Results

The following section is split by the simulation model’s severity and the real images test.

#### 4.2.1 Low Severity - Level 1

During the initial testing phase, the **MOANA** network was trained using dataset A1 and the simulation model’s severity level 1. The network’s performance was evaluated by altering the parameters, such as Batch Validation (BV) and the Number of Filters (NF). The results are shown for two scenarios, where BV is “Yes” or “No”, indicating its presence or absence. NF is 16 or 32, indicating the number of features used by the model. The aim was to determine the optimal parameters for better results. All the tests for severity level were conducted with 500 epochs.

The results, summarized in Table 2, highlight significant variations in model performance based on different parameter configurations. We verified that when BV was not implemented, and the NF was fixed at 16, our model’s SSIM values varied from 0.9473 to 0.9608 in different contrasts. Similarly, the NMI scores ranged from 0.7616 to 0.7704, and the corresponding NRMSE values from 0.1687 to 0.1938.

**Table 2.** Comparison of different model parameters and **MOANA** model results for SSIM, NMI, and NRMSE metrics, in dataset A1 and the simulation model’s severity level 1.

| Parameters |  |  | Results |  |  | Training Time |
| --- | --- | --- | --- | --- | --- | --- |
| BV | NF | Contrast | SSIM | NMI | NRMSE | (HH:MM:SS) |
| No | 16 | T1w | 0.9591 | 0.7616 | 0.1730 | 21:58:53 |
|  |  | T2w | 0.9608 | 0.7704 | 0.1687 |  |
|  |  | FLAIR | 0.9473 | 0.7638 | 0.1938 |  |
| Yes | 16 | T1w | 0.9682 | 0.7786 | 0.1527 | 22:06:22 |
|  |  | T2w | 0.9705 | 0.7811 | 0.1492 |  |
|  |  | FLAIR | 0.9593 | 0.7701 | 0.1639 |  |
| No | 32 | T1w | 0.9655 | 0.7756 | 0.1457 | 21:58:03 |
|  |  | T2w | 0.9654 | 0.7685 | 0.1480 |  |
|  |  | FLAIR | 0.9590 | 0.7818 | 0.1688 |  |
| Yes | 32 | T1w | 0.9745 | 0.7811 | 0.1497 | 22:07:03 |
|  |  | T2w | 0.9792 | 0.7821 | 0.1408 |  |
|  |  | FLAIR | 0.9803 | 0.8030 | 0.1580 |  |

However, we observed significant improvement in our model’s performance in all contrasts when we introduced BV. This is shown by higher SSIM values ranging from 0.9593 to 0.9705, indicating better structural similarity between predicted and ground truth images. Additionally, NMI values increased from 0.7786 to 0.7811, indicating enhanced capture of mutual information. NRMSE values decreased to a range of 0.1492 to 0.1639, signifying reduced error in model predictions.

Furthermore, increasing the NF from 16 to 32 without BV increased SSIM values in all the contrasts. A substantial boost in model performance is observed when combining an NF of 32 with BV, especially in SSIM values in different contrasts, particularly in FLAIR, which increased from 0.9590 to 0.9803. This denotes that a higher NF enables the model to extract more features and representations from the input images. At the same time, NMI values experienced a significant increase, from 0.7818 to 0.8030 in FLAIR contrast and the NRMSE values decreased from 0.1688 to 0.1580.

When using the BV technique in training **MOANA**’s correction model, the training time may increase slightly. The reason for this is that the validation process is performed for each batch, which takes some additional time. However, the increase in training time is not significant, for instance, in the case of dataset A1, the difference is only 0.6%. To complete 500 epochs, dataset A1 requires approximately 22 hours.

In Table 3, we present a comparison between MC2-Net, which includes the registration and the artifact correction networks, and **MOANA** models, in three different MRI contrasts: T1w, T2w, and FLAIR. All the tests were realized with 500 epochs.

**Table 3.** Comparison of the best SSIM and NMI metrics results in MC2-Net and **MOANA** models for three MRI contrasts.

| Model | Contrast | Metrics |  | Training Time<br>(Hours) |
| --- | --- | --- | --- | --- |
|  |  | SSIM | NMI |  |
| MC2-Net | T1w | 0.9891 | 0.8344 | 37+59* |
|  | T2w | 0.9787 | 0.7860 |  |
|  | FLAIR | 0.9726 | 0.7698 |  |
| **MOANA** | T1w | 0.9745 | 0.7811 | 22 |
|  | T2w | 0.9792 | 0.7821 |  |
|  | FLAIR | 0.9803 | 0.8030 |  |

\* Training time for registration and motion correction networks

When compared to MC2-Net, **MOANA** outperforms in T2w and FLAIR contrasts, with an increase of SSIM value from 0.9787 to 0.9792 and from 0.9726 to 0.9803, respectively. **MOANA** achieves higher SSIM values, indicating superior structural similarity between predicted and ground truth images, and higher NMI scores, suggesting better alignment between predicted and ground truth images in these two contrasts.

The training time varied across all experiments, depending on the model architecture and parameter configurations. MC2-Net had a substantially longer training time, averaging around 37 hours for the registration network (1000 epochs) and 59 hours for the correction network (500 epochs) [28]. In contrast, the **MOANA** model had significantly shorter training times, completing 500 epochs in approximately 22 hours. This reduction in training time with the **MOANA** model highlights its efficiency and computational feasibility for practical deployment in real-world scenarios.

Tripling the number of training images, as demonstrated in Table 4, suggests that using a larger and more diverse dataset enhances model performance. This expanded dataset likely improves the model’s ability to generalize, leading to more accurate predictions. Consequently, the model was trained for a total of 1000 epochs due to the increase in dataset size.

**Table 4.** Comparison of different model parameters and **MOANA** model results for SSIM, NMI, and NRMSE metrics, in dataset B1 and the simulation model’s severity level 1.

| Parameters |  |  | Results |  |  | Training Time<br>(D-HH:MM:SS) |
| --- | --- | --- | --- | --- | --- | --- |
| BV | NF | Contrast | SSIM | NMI | NRMSE |  |
| No | 16 | T1w | 0.9551 | 0.7613 | 0.1394 | 3-11:59:25 |
|  |  | T1CE | 0.9617 | 0.7775 | 0.1609 |  |
|  |  | T2w | 0.9573 | 0.7721 | 0.2271 |  |
|  |  | FLAIR | 0.9455 | 0.7499 | 0.1983 |  |
| Yes | 16 | T1w | 0.9574 | 0.7642 | 0.1365 | 3-12:24:43 |
|  |  | T1CE | 0.9624 | 0.7798 | 0.1570 |  |
|  |  | T2w | 0.9605 | 0.7743 | 0.1938 |  |
|  |  | FLAIR | 0.9647 | 0.7975 | 0.1559 |  |
| No | 32 | T1w | 0.9601 | 0.7658 | 0.1346 | 3-12:08:32 |
|  |  | T1CE | 0.9674 | 0.7825 | 0.1711 |  |
|  |  | T2w | 0.9650 | 0.7849 | 0.1765 |  |
|  |  | FLAIR | 0.9709 | 0.7942 | 0.1628 |  |
| Yes | 32 | T1w | **0.9682** | 0.7843 | 0.1427 | 3-12:25:53 |
|  |  | T1CE | **0.9749** | 0.7914 | 0.1420 |  |
|  |  | T2w | **0.9715** | 0.7927 | 0.1421 |  |
|  |  | FLAIR | **0.9771** | 0.7968 | 0.1313 |  |

The findings indicate that using BV generally enhances model performance across all contrast types. For instance, in FLAIR images with 16 filters, the SSIM improves from 0.9455 (without BV) to 0.9647 with BV, while the NRMSE decreases from 0.1983 to 0.1559, reflecting more precise image reconstructions. Additionally, increasing the NF from 16 to 32 boosts model performance, even without BV. For example, in T1w images without BV, the SSIM increases from 0.9551 to 0.9601, and the NRMSE reduces from 0.1394 to 0.1346. When BV is used with 32 filters, the improvements are even more significant. In T1w images, for example, the SSIM rises to 0.9682, and the NRMSE falls to 0.1427. The most optimal results are achieved when both BV and 32 filters are applied. In this setup, the highest SSIM value is observed in FLAIR images (0.9771), and the lowest NRMSE (0.1313) indicates the most accurate artifact correction.

Despite these performance improvements, training times across different configurations remain similar, with all models completing in approximately 3 days and 12 hours. This indicates that the application of BV and increasing the NF does not considerably impact computational costs. Thus, both strategies enhance model performance, with the combination of BV and 32 filters yielding the best results across all contrast types.

Figure 5 shows examples of the output of our correction model after the input of corrupted images with severity level 1. As can be observed, the SSIM values of motion-corrupted images are significantly lower than the SSIM values of motion-corrected ones.

**Figure 5.**
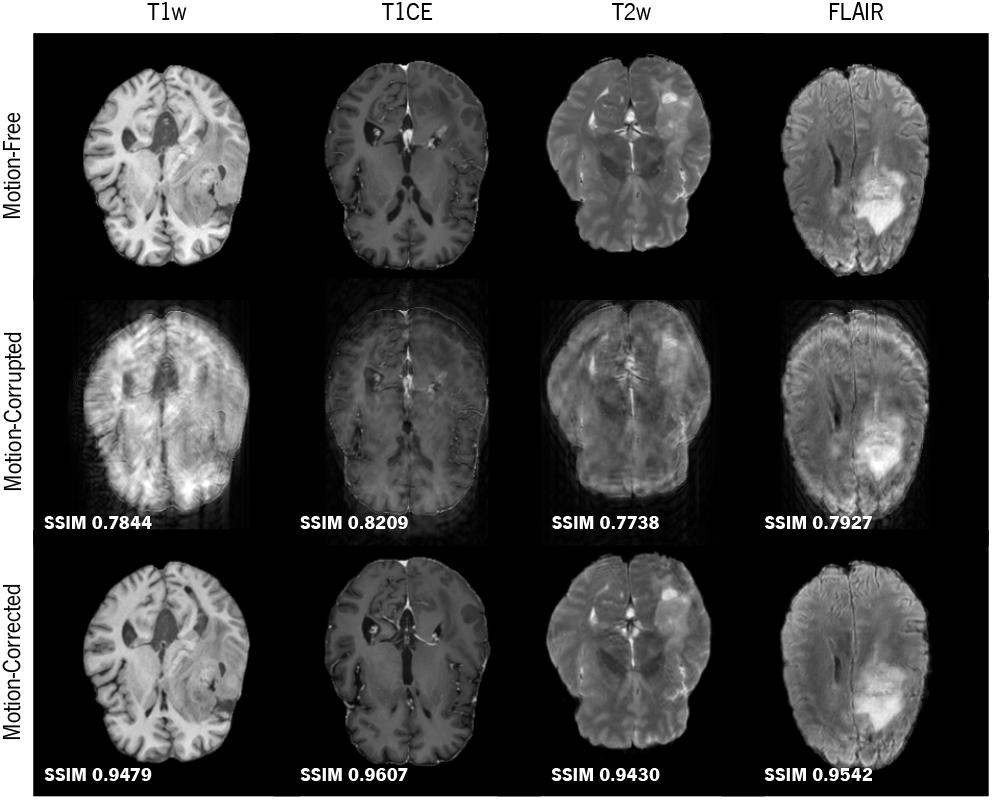
Example images of correction in dataset B1 and the simulation model’s severity level 1.

The average SSIM of the output images from motion-free input images was calculated to be 0.9998, 0.9996, and 0.9997 for T1w, T2w, and FLAIR contrasts, respectively. To test the effectiveness of the model in correcting artifacts without adding new details, we conducted another test. T1w input images were motion-corrupted, while T2w and FLAIR input images were motion-free. In this scenario, the MC2-Net achieved an SSIM of 0.9905, 0.9821, and 0.9746 for T1w, T2w, and FLAIR contrasts, respectively. Moreover, the **MOANA** model achieved values of SSIM equal to 0.9969, 0.9899, and 0.9908 for the same contrasts. These results demonstrate that the model can efficiently correct motion artifacts and identify motion-free images without modifying their information.

#### 4.2.2 High Severity - Level 2

Table 5 compares the different model parameters and their respective performance metrics when training **MOANA** using the A1 dataset with the severity level 2. The experiments were performed over 500 epochs.

**Table 5.** Comparison of different model parameters and **MOANA** model results for SSIM, NMI, and NRMSE metrics, in dataset A1 and the simulation model’s severity level 2.

| Parameters |  |  | Results |  |  | Training Time |
| --- | --- | --- | --- | --- | --- | --- |
| BV | NF | Contrast | SSIM | NMI | NRMSE | (HH:MM:SS) |
| No | 16 | T1w | 0.9218 | 0.6648 | 0.2515 | 21:51:36 |
|  |  | T2w | 0.9414 | 0.7376 | 0.2238 |  |
|  |  | FLAIR | 0.9610 | 0.7661 | 0.1393 |  |
| Yes | 16 | T1w | 0.9243 | 0.6777 | 0.2309 | 22:05:07 |
|  |  | T2w | 0.9428 | 0.7411 | 0.2105 |  |
|  |  | FLAIR | 0.9624 | 0.7718 | 0.1368 |  |
| No | 32 | T1w | 0.9387 | 0.7364 | 0.2422 | 21:51:10 |
|  |  | T2w | 0.9489 | 0.7506 | 0.1925 |  |
|  |  | FLAIR | 0.9635 | 0.7747 | 0.1323 |  |
| Yes | 32 | T1w | 0.9425 | 0.7498 | 0.2308 | 22:06:42 |
|  |  | T2w | 0.9543 | 0.7667 | 0.1536 |  |
|  |  | FLAIR | 0.9637 | 0.7779 | 0.1292 |  |

The results showed that when BV was omitted and NF was set to 16, the model’s SSIM values ranged from 0.9218 to 0.9610, NMI scores between 0.6648 to 0.7661, and NRMSE values from 0.1393 to 0.2515. However, the introduction of BV significantly improved the model’s performance in all contrasts, yielding higher SSIM values (ranging from 0.9243 to 0.9624), increased NMI scores (ranging from 0.6777 to 0.7718), and reduced NRMSE values (up to 8% of reduction).

Furthermore, the increase in NF from 16 to 32 enhanced the model’s performance, particularly when used alongside BV. For instance, in the T2w contrast, when NF was increased from 16 to 32 and BV was applied, the SSIM value rose from 0.9428 to 0.9543, representing a clear improvement in the structural similarity between predicted and ground truth images. Analogously, the NMI score increased from 0.7411 to 0.7667 and the NRMSE value decreased from 0.2105 to 0.1536. This illustrates how the changes in NF and BV allowed the model to capture more detailed information from the input images, resulting in clearer and more accurate predictions. It is worth noting that the model requires approximately 22 hours to complete 500 epochs.

Table 6 shows a comparison of the results obtained by using different model parameters with the **MOANA** model in four image contrasts, for the A2 dataset at severity level 2 of the simulation model. The tests were conducted during 500 epochs.

**Table 6.** Comparison of different model parameters and **MOANA** model results for SSIM, NMI, and NRMSE metrics, in dataset A2 and the simulation model’s severity level 2.

| Parameters |  |  | Results |  |  | Training Time |
| --- | --- | --- | --- | --- | --- | --- |
| BV | NF | Contrast | SSIM | NMI | NRMSE | (D-HH:MM:SS) |
| No | 16 | T1w | 0.8665 | 0.6349 | 0.3860 | 1-04:03:09 |
|  |  | T1CE | 0.9367 | 0.7225 | 0.2330 |  |
|  |  | T2w | 0.8606 | 0.6365 | 0.3453 |  |
|  |  | FLAIR | 0.9604 | 0.7561 | 0.1408 |  |
| Yes | 16 | T1w | 0.9220 | 0.6761 | 0.2483 | 1-04:27:09 |
|  |  | T1CE | 0.9383 | 0.7209 | 0.2301 |  |
|  |  | T2w | 0.9421 | 0.7353 | 0.2342 |  |
|  |  | FLAIR | 0.9613 | 0.7643 | 0.1433 |  |
| No | 32 | T1w | 0.9533 | 0.7690 | 0.2334 | 1-03:52:22 |
|  |  | T1CE | 0.9344 | 0.7263 | 0.2691 |  |
|  |  | T2w | 0.9377 | 0.7282 | 0.2655 |  |
|  |  | FLAIR | 0.9545 | 0.7692 | 0.2273 |  |
| Yes | 32 | T1w | 0.9542 | 0.7679 | 0.1517 | 1-04:10:43 |
|  |  | T1CE | 0.9464 | 0.7329 | 0.2255 |  |
|  |  | T2w | 0.9482 | 0.7374 | 0.2321 |  |
|  |  | FLAIR | 0.9664 | 0.7784 | 0.1350 |  |

By analyzing the results in Table 6, it is perceptible that when BV is not applied, and the NF is set to 16, the SSIM scores range between 0.8606 and 0.9604 for all four contrasts. With BV applied and NF set to 16, the SSIM increased to up to 0.9613. On the other hand, by increasing the NF to 32 without applying BV, the results are superior in some metrics. For example, for the FLAIR contrast, without BV and with an NF of 32, the NRMSE is 0.2273. However, when shifting BV to the “Yes” status with an NF of 32, there is a significant reduction in NRMSE to 0.1350, indicating a considerable improvement in the model’s accuracy.

In addition, the presence of BV can positively affect the metrics in other contrasts. For instance, for the T1CE contrast, activating BV with an NF of 32 increases SSIM from 0.9344 to 0.9464, which shows an improvement in the structural similarity of the images processed by the model. Regarding the training time for 500 epochs, the process was completed in 28 hours.

By tripling the number of training images, the results presented in Table 7 show that by having access to a more extensive and diverse dataset for learning, the model performs better. This increased dataset potentially enhances the model’s ability to generalize and produce more accurate predictions. Due to the increased dataset size, the training endured for 1000 epochs.

**Table 7.** Comparison of different model parameters and **MOANA** model results for SSIM, NMI, and NRMSE metrics, in dataset B1 and the simulation model’s severity level 2.

| Parameters |  |  | Results |  |  | Training Time |
| --- | --- | --- | --- | --- | --- | --- |
| BV | NF | Contrast | SSIM | NMI | NRMSE | (D-HH:MM:SS) |
| No | 16 | T1w | 0.9383 | 0.7387 | 0.1571 | 3-11:59:07 |
|  |  | T1CE | 0.9451 | 0.7664 | 0.1958 |  |
|  |  | T2w | 0.9402 | 0.7564 | 0.2740 |  |
|  |  | FLAIR | 0.9313 | 0.7318 | 0.2233 |  |
| Yes | 16 | T1w | 0.9395 | 0.7441 | 0.1577 | 3-12:49:35 |
|  |  | T1CE | 0.9453 | 0.7666 | 0.1838 |  |
|  |  | T2w | 0.9452 | 0.7603 | 0.2228 |  |
|  |  | FLAIR | 0.9530 | 0.7876 | 0.1759 |  |
| No | 32 | T1w | 0.9399 | 0.7402 | 0.1588 | 3-12:11:21 |
|  |  | T1CE | 0.9482 | 0.7596 | 0.1968 |  |
|  |  | T2w | 0.9488 | 0.7699 | 0.1970 |  |
|  |  | FLAIR | 0.9542 | 0.7808 | 0.1808 |  |
| Yes | 32 | T1w | 0.9479 | 0.7672 | 0.1290 | 3-12:55:43 |
|  |  | T1CE | 0.9607 | 0.7929 | 0.1695 |  |
|  |  | T2w | 0.9530 | 0.7708 | 0.1452 |  |
|  |  | FLAIR | 0.9647 | 0.7966 | 0.1506 |  |

The results indicate that the model consistently performs better when BV is used in all contrasts. For instance, in FLAIR images with 16 filters, SSIM increased from 0.9313 to 0.9530 with BV, while NMI rose from 0.7318 to 0.7876.

Moreover, when examining the influence of NF, we see that increasing NF generally leads to better results. This tendency is evident in T2w images, where SSIM increased from 0.9402 to 0.9488 when going from 16 to 32 filters without BV. Conversely, the effect of NF is more pronounced when BV is used. This is exemplified in T1CE images, where SSIM improved from 0.9453 to 0.9607 when BV was used with 32 filters. The model was trained for approximately 94 hours.

#### 4.2.3 Real Images

First, the **MOANA** correction model was trained with simulated images due to the difficulty of obtaining motion-free and motion-corrupted images of the same slice. Then, the model was inferred with data from four subjects (dataset C1) containing real movement. The correction results for the T1CE contrast are presented in Figure 6. Even when applied to real images, the figure shows that our correction model successfully corrected and removed movement from the images. This validates the method’s effectiveness and applicability in real-world scenarios, providing a reliable solution for correcting images affected by motion.

**Figure 6.**
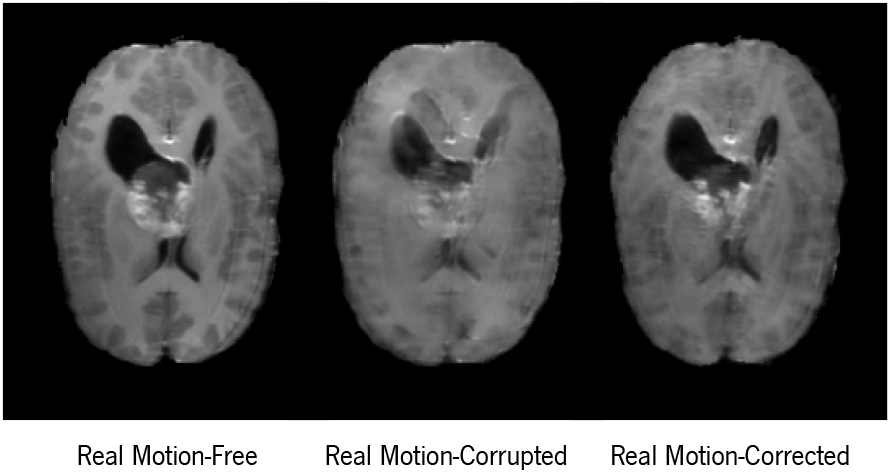
Example images of motion correction in the contrast T1CE of dataset C1, containing real motion-free, motion-corrupted and motion-corrected images.

### 4.3 Limitations

The scarcity of real images that contain artifacts and the corresponding ground truth images is bridged by **MOANA**’s simulation model. As such, by following a retrospective approach, **MOANA** requires both the source image and the corrupted image to apply the correction process accurately. With this, the inference of real images in such a model depends directly on the availability of corruption-free slices from the same subject. This is a general issue that should be addressed by the scientific community by gathering non-curated and artifact-corrupted scans and making them publicly available for research use.

Additionally, our correction model was able to attain an average SSIM for all contrasts over 0.93. In cases where the artifacts were subtle, the correction using 32 filters and BV showed noteworthy improvements when compared with the use of 16 filters and no BV. However, we believe that these results could be refined by resorting to distributed settings, as we defer this to future work.

## 5 Conclusion

We propose **MOANA**, an end-to-end pipeline for artifact correction in MRI images. By offering two models, **MOANA** is able to efficiently mimic common artifacts in previously curated data and further correct them. **MOANA**’s capability to handle artifacts in various contrasts, combined with its efficiency in correcting different levels of distortion, particularly severe artifacts, makes it an advantageous tool in medical imaging analysis. Although there are still challenges in correcting subtle artifacts, the model’s performance exhibits improvements over existing methods, indicating a promising direction for future research and development in the field of artifact correction in MRI. With continued advancements, **MOANA** has the potential to assist clinicians in obtaining clearer and more trustworthy diagnoses.

